# A dramatic impairment of the antitumor activity of human Vγ9Vδ2 T cells is induced by TGF-β through significant phenotype, transcriptomic and metabolic changes

**DOI:** 10.1101/2022.09.22.509026

**Authors:** Chirine Rafia, Clément Loizeau, Ophélie Renoult, Christelle Harly, Claire Pecqueur, Noémie Joalland, Emmanuel Scotet

## Abstract

Despite significant advances, the eradication of cancer remains a clinical challenge which justifies the urgent exploration of additional therapeutic strategies such as immunotherapies. Human peripheral Vγ9Vδ2 T cells represent an attractive candidate subset for designing safe, feasible and effective adoptive T cell transfer-based therapies. However, following their infiltration within tumors, γδ T cells are exposed to various regulating constituents and signals from the tumor microenvironment (TME), which severely alter their antitumor functions. Here, we show that TGF-β, whose elevated production in some solid tumors is linked to a poor prognosis, interferes with the antigenic activation of human Vγ9Vδ2 T cells *in vitro*. This regulatory cytokine strongly impairs their cytolytic activity, which is accompanied by the induction of particular phenotypic, transcriptomic and metabolic changes. Collectively, these observations provide informations for better understanding and targeting the impact of TME components to regulate the antitumor activity of human T cell effectors.

## Introduction

In the wake of revolutionary cancer immunotherapies, adoptive transfer strategies harnessing effector functions of particular immune cell subsets have been proposed. Among the most attractive candidates, human peripheral Vγ9Vδ2 T lymphocytes, which represent 1 to 10% of blood CD3+ T lymphocytes in healthy adults, contribute to host defenses against various infections and malignancies^1,2^. These conserved and non-alloreactive unconventional T lymphocytes are selectively activated in a TCR/CD3-dependent but MHC-independent, manner by non-peptidic small metabolites of isoprenoid synthesis pathways called PAg (*phosphoantigen*), such as IPP (*isopentenyl pyrophosphate*). This peculiar and exquisitely specific stress sensing process, which remains ill defined, also involves BTN (*butyrophilin*)2A/BTN3A, which are expressed at the surface of target cells ^3–5^. The sensitization of target cells with elevated pinocytic activity by pharmacological NBP (*amino-bisphosphonate*) compounds, such as Zoledronate, which inhibit a key IPP-degrading enzyme of the mammalian mevalonate pathway, upregulate the reactivity of Vγ9Vδ2 T lymphocytes^6^. This *Self*-targeted reactivity process needs to be tightly controlled by a set of molecules, such as NK (*natural killer*) receptors, adhesion molecules, Toll-*like* receptors or immune checkpoint inhibitors. The activating NKG2D (*natural killer group 2, member D*) receptor is expressed by cytotoxic immune cell subsets, including Vγ9Vδ2 T lymphocytes. Once engaged by stress-induced determinants that are barely, if not, expressed by healthy cells, these receptors play a key role in sustaining TCR/CD3-induced reactivities and subsequent antitumor functions, such as perforin/granzyme-mediated cytolysis. Vγ9Vδ2 T cells integrate activation and inhibition signals to deliver, or not, rapid and strong functional responses, such as proliferation, cytolysis and cytokines (i.e. TNF (*tumor necrosis factor*)-α, IFN (*interferon*)-γ) release^7^.

Importantly, Vγ9Vδ2 T lymphocytes sense and eliminate a broad range of human circulating and solid tumor cells. The range of tumor cell lines detected by Vγ9Vδ2 T cells, which was initially restricted to circulating tumors (e.g. hematopoietic tumors), has been next extended to various solid tumors (e.g. renal cancer and colon carcinomas)^8^. The development of clinical grade synthetic agonist molecules such as NBP, PAg or anti-BTN3A agonist antibodies enabled the implementation of passive or active Vγ9Vδ2 T cell immunotherapies with modest therapeutic effects^9^. However, while several groups initially reported tumor infiltration by human γδ T cells, additional work evidenced that some tumor-associated γδ T cells display pro-tumor functions^10,11^. This result highlights the role played by the TME (*tumor microenvironment*) on tumor-infiltrated γδ T cells in fostering immune escape. Among TME factors, TGF-β (*transforming growth factor-β*), a highly conserved multi-functional cytokine, is secreted by different cell types (e.g. lymphocytes, macrophages) and regulates various biological functions of tumor and T cells^12,13^. Indeed, different studies have shown its strong suppressive activity within solid tumors by inducing lymphocyte cell death or converting effector T cells into T helper 17 or T regulatory lymphocytes (see ^12^ for a review).

On the basis of the strong and well described impact of TGF-β on antitumor αβ T cells reactivities, we investigated its influence on the human effector Vγ9Vδ2 T cell subset. Our results evidenced that antitumor functions of Vγ9Vδ2 T cells are greatly impaired by TGF-β. More importantly, they demonstrate that this impairment is not only profound, as it affects the transcriptome and the metabolism of Vγ9Vδ2 T lymphocytes, but also long lasting.

## Materials & Methods

### Reagents, human Vγ9Vδ2 T and tumor cells

Human Vγ9Vδ2 T lymphocytes were amplified from PBL (*peripheral blood lymphocytes*) obtained from healthy donor blood samples provided by the Etablissement Français du Sang (EFS, Nantes, France) and after Ficoll density centrifugation (Eurobio). For specific *ex vivo* expansions of peripheral allogeneic human Vγ9Vδ2 T lymphocytes, PBL were incubated with 3 μM of BrHPP (bromohydrin pyrophosphate), kindly provided by Innate Pharma (Marseille, France) in RPMI medium supplemented with 10% heat-inactivated fetal calf serum, 2 mM L-glutamine, 10 mg/mL streptomycin, 100 IU/mL penicillin, and 100 IU/mL human recombinant IL-2 (Proleukin, Novartis). After 4 days, cell cultures were supplemented with 300 IU/mL IL-2. On day 21, the purity of cultures was checked by flow cytometry and the population with a purity >90% were conserved. Human Raji (Burkitt’s lymphoma) and SKOV3 (ovarian carcinoma) were cultured in complete RMPI (Roswell Park Memorial Institute, Gibco) and DMEM (Dulbecco’s Modified Eagle’s Medium, Gibco) medium, supplemented with 10% heat-inactivated fetal calf serum (Gibco), 2 mM L-glutamine (Gibco), 10 mg/mL streptomycin (Gibco), 100 IU/mL penicillin (Gibco), respectively. Human primary GBM (Glioblastoma Multiforme) cultures were obtained after mechanical dissociation of surgical resection tumor samples from patients. All procedures involving human patients were performed in accordance with the ethical standards of the ethic national research committee and with the 1964 Helsinki declaration and its later amendments or comparable ethical standards. Informed consent was obtained from all individual patients included in this study. Primary cultures were established and stored at −180°C. Cell-frozen vials were grown in defined medium (DMEM/Ham F12 (Gibco), 2 mmol/L l-glutamine (Gibco), N2- and B27-supplement (Gibco), 2 μg/mL heparin (Sigma Aldrich), 20 ng/mL EGF and 25 ng/mL bFGF (Peprotech), and 100 IU/mL penicillin and 100 mg/mL streptomycin (Gibco)) at 37°C in a humidified atmosphere with 5% CO_2_. All experiments were performed with GBM cells in culture for less than 3 months and cells were routinely checked for mycoplasma contamination. Recombinant human transforming growth factor β1 (premium grade, Miltenyi Biotec) was diluted and used at the indicated concentrations.

### Flow cytometry

Human Vγ9Vδ2 T lymphocytes were stained with FITC-labeled anti-human TCR Vδ2 mAb (#IMMU389, Beckman Coulter) and APC-labeled anti-human CD3ε mAb (#UCHT1, Beckman Coulter), to check the purity of the populations. Human Vγ9Vδ2 T lymphocytes were phenotyped using PE-labeled anti-NKG2A (#Z199, Beckman Coulter), PE-labeled anti-NKG2D (#ON72, Beckman Coulter), PE-labeled anti-CD85j/ILT2 (#HP-F1 Beckman Coulter), APC-labeled anti-ICAM1 (#HA58, Biolegend), Acquisition was performed using an Accuri C6 PLUS flow cytometer (BD Biosciences) and the collected events were analyzed using FlowJo software (Treestar).

### Functional assays *in vitro*

For CD107a surface mobilization assays, primary human GBM cells or cell lines were sensitized, or not, with various quantities of zoledronate (Zometa, Sigma) overnight before incubation with human Vγ9Vδ2 T lymphocytes (effector to target ratio 1:1). Cocultures were performed in RPMI medium containing 5 μM monensin (Sigma) and Alexa Fluor 647-labeled anti-human CD107a mAb (#H4A3, Biolegend). After 4 h, human Vγ9Vδ2 T lymphocytes were collected and stained with a FITC-labeled anti-human TCR Vδ2 mAb (#IMMU389, Beckman Coulter) and analyzed by flow cytometry. To perform cytotoxic assays, tumor cells were previously treated with Zol overnight and then incubated with ^51^Cr (75 μCi/10^6^ cells) for 1 h. After washes, cells were cocultured with human Vγ9Vδ2 T lymphocytes (effector to target ratio 10:1) for 4 h. The activity of ^51^Cr released in supernatants was measured using a MicroBeta counter (Perkin Elmer). Specific target cell lysis was calculated as follows: % Lysis=((Experimental release-Spontaneous release)/(Maximum release-Spontaneous release)) x 100. Spontaneous and maximum release values were obtained by adding culture medium or 1% Triton X−100, respectively, to ^51^Cr-labeled target cells in the absence of human Vγ9Vδ2 T lymphocytes.

### Transcriptional analyses

Briefly, RNA from dry pellets of Vγ9Vδ2 T lymphocytes was extracted and purified using the NucleoSpin RNA plus isolation kit from Macherey-Nagel. DGE-sequencing was performed by the BIRD core facility and data were analyzed using R programming languages and tools.

### Metabolic analyses

Mitochondrial oxygen consumption (OCR) and extracellular acidification rate (ECAR) were measured in non-buffered medium containing 0.2 mM cystine supplemented with glucose (5 mM), pyruvate (1 mM), and glutamine (2 mM) using an XF24 Analyzer (SeaHorse Bioscience). Briefly, Vγ9Vδ2 T lymphocytes are washed in the analysis medium and plated at 8 × 10^5^ cells per well using Cell-Tak (Corning) at 22,4 μg/mL. The plate is then placed at 37°C to equilibrate for 1h prior to starting the analysis. First, points of measurement are taken every 10 minutes. After 3 cycles, BrHPP is next injected and 3 measures are taken rapidly, every 3 min, followed by 5 cycles of 10 min. PMA+ionomycin is then injected and the same protocol of measurement is applied than after BrHPP. Specific mitochondrial respiration in the presence of a stimulus (BrHPP, PMA-ionomycin) was calculated as follows: difference between the mean of the 3 values of OCR in the absence of substrate and the mean of the 4 values of OCR after injection of the substrate. Analysis of the obtained values were performed using Wave 2.6 (Agilent) software.

### Statistical analyses

Data are expressed as mean ± SD and were analyzed using GraphPad Prism 6.0 software (GraphPad Software). Two-way ANOVA and Mann & Whitney tests were calculated to assess or not for significant differences.

## Results

### Elevated levels of TGF-β transcripts are detected in human brain tumors with a poor prognosis

The abundance of TGF-β transcripts was analyzed within human tumors, such as brain tumors, using GEPIA (*Gene Expression Profiling Interactive Analysis*, http://gepia.cancer-pku.cn/) which is an interactive web-based tool developed to deliver fast and customizable functionalities based on TCGA (*The Cancer Genome Atlas*) and GTEx (*Genotype-Tissue Expression*) data^14^. These *in silico* transcriptomic analyses evidenced high quantities of TGF-β transcripts in Glioblastoma (GBM) and Low-Grade Gliomas (LGG), as compared to normal brain tissues (***Figure 1A*** and ***1B***). GEPIA also provides interactive functions such as correlation and patient survival analyses. Interestingly, these analyses revealed that increased abundance of TGF-β transcripts in brain tumors is correlated to a dismal prognosis (i.e. overall and disease-free survivals) of these brain tumor patients (***Figure 1C***). These analyses indicate that elevated levels of TGF-β transcripts are produced in human brain cancers with poor prognosis.

**Figure 1.**
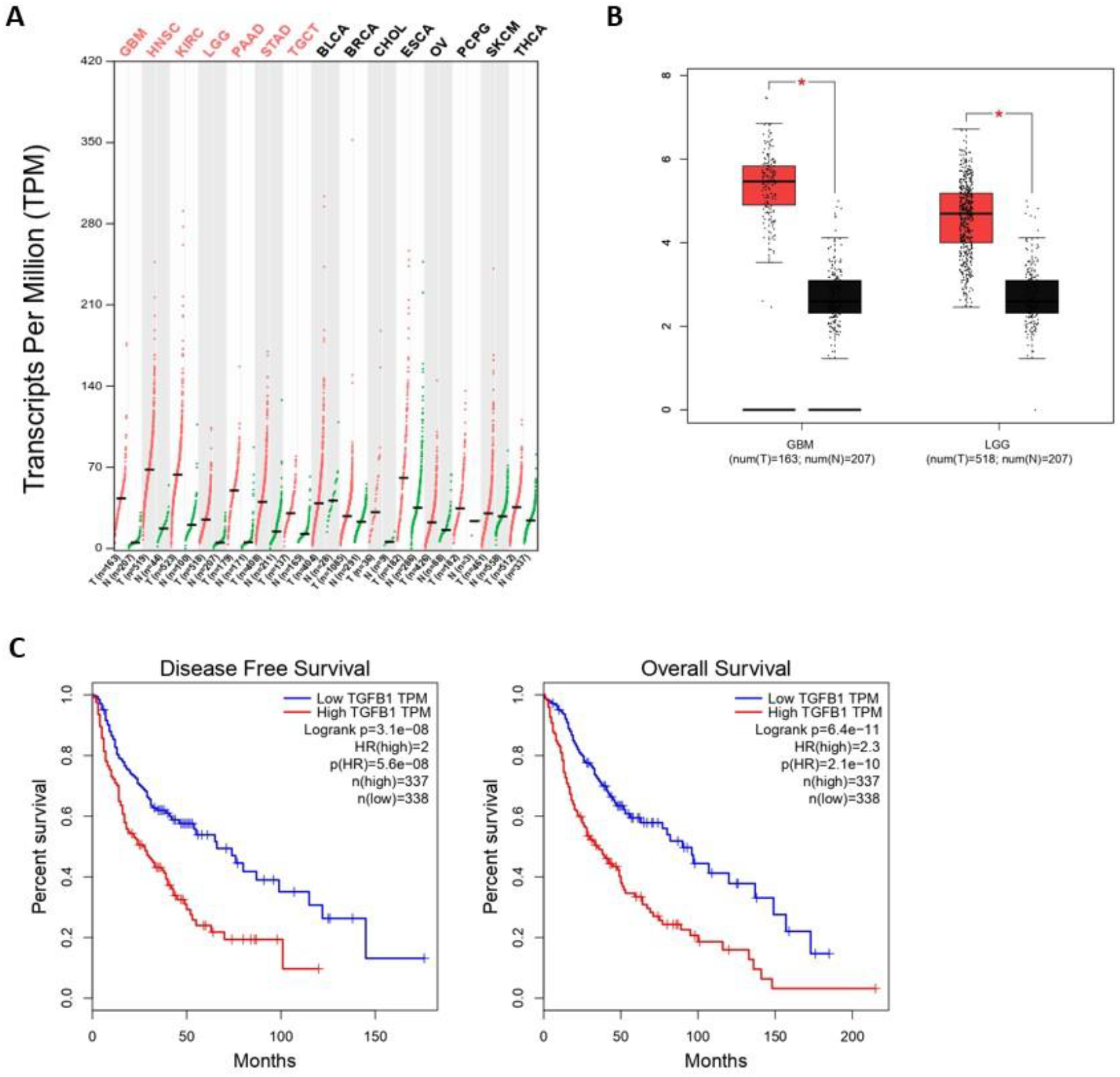
High levels of TGF-β transcripts are present in human brain tumors with a poor prognosis. **(A)** The abundance of human TGF-β transcripts (TPM, transcripts per million) across indicated various cancers (T) and the normal tissues associated (N) was analyzed and compared by matching TCGA and GTEx data. (B) Comparative analyses on human brain tumors (GBM, Glioblastoma; LGG, Low Grade Glioma) ^*^p<0,05. **(C)** Survival curves (Disease free, *left*; Overall, *right*) for levels (low, *blue*; high, *red*) of TGFB1 transcripts analyzed for GBM and LGG (Kaplan-Meier). Data were obtained from GEPIA (http://gepia.cancer-pku.cn/).

### TGF-β inhibits the antitumor cytotoxic reactivity of human Vγ9Vδ2 T cells

Considering elevated levels of TGF-β transcripts in brain human tumors with a dismal prognosis and the role of Vγ9Vδ2 T cells against GBM cells^15,16^, we investigated the inhibitory effect induced by TGF-β. ^51^Cr-release assays performed using Raji and GBM cells (primary culture) as targets showed that zoledronate-induced cytolytic activities were inhibited in the presence of TGF-β (***Figure 2A***). Vγ9Vδ2 T cells which have been pre-incubated for 3 days with TGF-β (10 ng/mL) expressed significantly reduced levels of T cell surface-translocated CD107a when activated by increasing doses of anti-CD3 antibody, as compared to control conditions (*untreated*). Importantly, both natural and zoledronate induced-reactivities (CD107a) of TGF-β-treated Vγ9Vδ2 T cells against tumor cells from ovarian cancer (SKOV3) and GBM primary cultures (GBM-1), were significantly reduced, as compared to control conditions (*untreated*) (***Figure 2B***). Interestingly, strong antigenic activation conditions (zoledronate, 10 μM) reduced the intensity of this inhibition.

**Figure 2.**
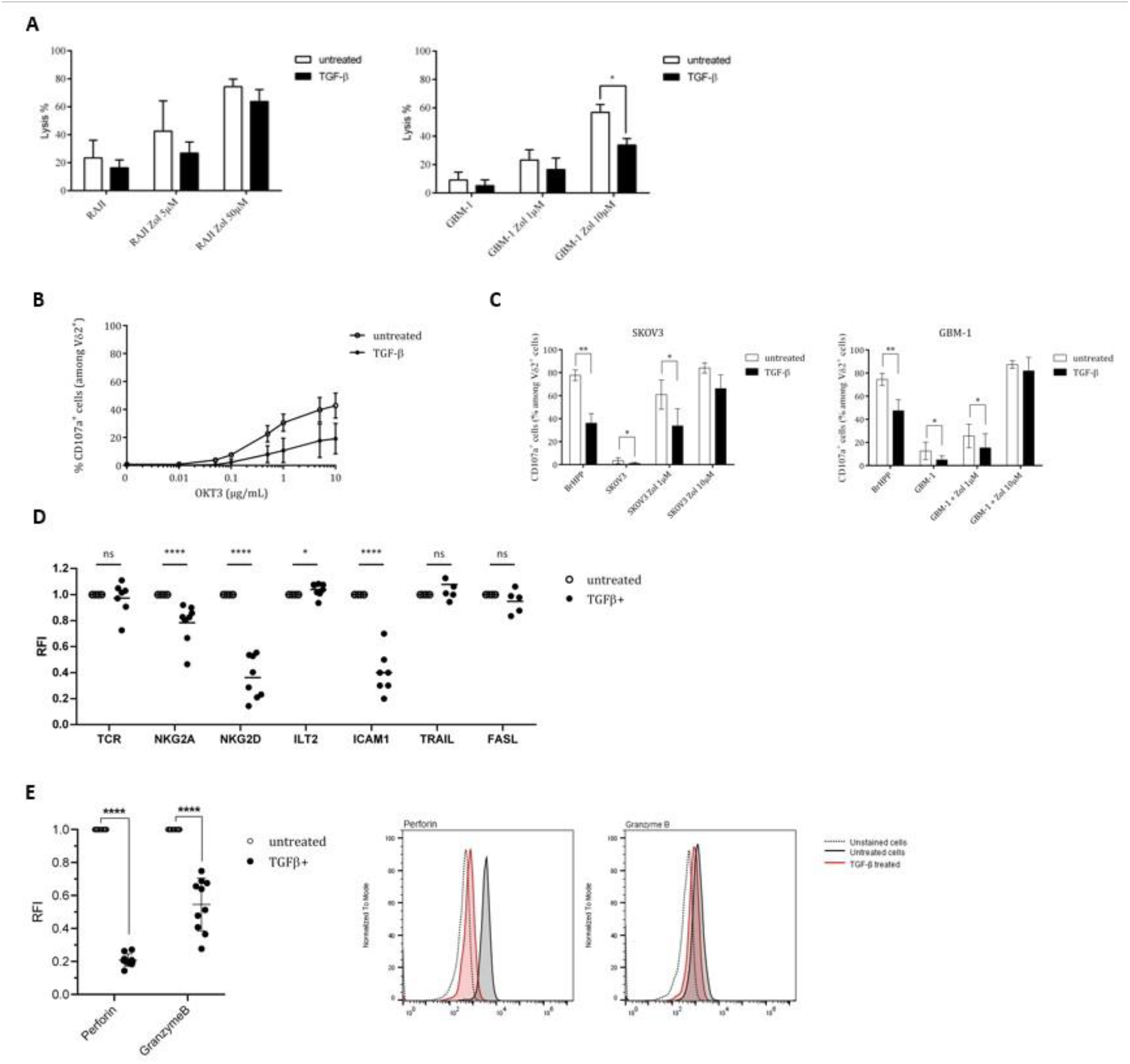
TGF-β inhibits both antitumor cytotoxicity and antigenic activation of human Vγ9Vδ2 T cells. PBL-derived Vγ9Vδ2 T cells were pretreated or not with TGF-β (10 ng/mL) for 3 days. **(A)** Anti-tumor cytotoxic reactivities of TGF-β-treated (*closed*) *vs* untreated (*open*) Vγ9Vδ2 T lymphocytes against human tumor cells (Raji, left; GBM-1, right) using ^51^Cr release assays. Target cells have been pretreated, or not, with zoledronate (Zol) at 5 and 50 μM. Results are expressed as % of cytotoxicity (mean ± SD). **(B)** and **(C)** Frequency of CD107a^+^ cells among TGF-β-treated (*closed*) *vs* untreated (*open*) TCRVδ2^+^ lymphocytes (%) after stimulation with increasing concentrations of anti-CD3ε mAb (OKT3) **(B)** or zoledronate (Zol)-treated, or not, human SKOV3 (*left*) or GBM-1 (*right*) tumor cells **(C)**. E/T ratio 1. Positive control: BrHPP (3 μM). n≥5, mean ± SD. **(D)** and **(E)** Expression levels of activation-related molecules (TCR, NKG2A, NKG2D, ICAM1, TRAIL, FASL) **(D)** and cytolytic (Perforin, Granzyme B) molecules **(E)** by TGF-β-treated (*closed*) *vs* untreated (*open*) TCRVδ2^+^ lymphocytes measured by flow cytometry. RFI: Relative Fluorescence Intensity. Mann-Whitney ^*^p<0,05 ; ^**^p<0,01 ; ^****^p<0,001.

In an effort to provide mechanistic explanation(s) for this diminished activation potential and antitumor reactivity, the levels of expression of key selected surface molecules were analyzed and compared by flow cytometry with a focus on markers playing a role in the antitumor reactivity such as TCR, ICAM-1, activatory (NKG2D) or inhibitory (CD94/NKG2A; ILT2) NK receptors as well as cytolytic pathways associated molecules (perforin (perf), granzyme B (GmzB), Fas Ligand (FASL), TRAIL)) (***Figure 2C***). The most striking effect of this TGF-β exposure was the strongly reduced expression of the activating receptor NKG2D, the cell adhesion molecule ICAM-1 and perforin/granzyme B which represent key contributors of antitumor cytolytic pathways used by Vγ9Vδ2 T cells. TGF-β exposure did not significantly affect the expression of TCR, FASL and TRAIL molecules and slightly affected the expression of inhibitory receptors. Following dose-responses analyses to determine optimal TGF-β concentrations for inducing a significant reduction of NKG2D expression (***Supplemental Figure S1A***), the kinetics of effects induced by TGF-β were established, at the protein level, by measuring the levels of expression of NKG2D at different time-points following addition of TGF-β (***Supplemental Figure S1B***). The expression of NKG2D remained stable for about 4 h in the presence of TGF-β and started to decrease after 8 h by reaching its lowest expression levels between 24 h and 72 h. Interestingly, next experiments showed that, following exposure to TGF-β for different durations and washes, the impact of this cytokine on NKG2D expression was lasting for days, with a partial restoration of initial expression levels 5 days after withdrawing TGF-β (***Supplemental Figure S1C***).

The intracellular levels of expression of cytolytic molecules from TGF-β-treated and untreated Vγ9Vδ2 T cells were measured and compared (***Figure 2D***). These analyses focused on perforin/granzyme B, which constitute the major pathway contributing to the antitumor cytolytic activity of Vγ9Vδ2 T cells. TGF-β-treated Vγ9Vδ2 T cells expressed significantly lower levels of perforin and granzyme B, as compared to untreated conditions (*Relative Fluorescence Intensity* (RFI)=20 % and 50 %, respectively). Taken together, these results show that TGF-β significantly inhibits the antigenic reactivity as well as the antitumor cytolytic potential and activity of human Vγ9Vδ2 T cells.

### The transcriptome of human Vγ9Vδ2 T cells is significantly modified upon TGF-β exposure

To check whether inhibition of the antitumor Vγ9Vδ2 T cells reactivity induced by TGF-β could also be linked to transcriptome modifications, comparative DGE-seq analyses from bulk RNA-seq data have been performed. RNA samples were prepared from PBL-derived Vγ9Vδ2 T cells exposed, or not, for 3 days to TGF-β *in vitro*. 89 genes (40 upregulated, 49 downregulated) were found differentially expressed (fold change > 2, adjusted *p* value < 0.05) between TGF-β-treated and - untreated groups (***Figure 3A***). Interestingly, MA plot analysis, which represents log fold-change versus mean expression of genes between TGF-β treated and non-treated Vγ9Vδ2 T cells, shows that well-expressed down-regulated genes mainly encode for surface-bound receptors such as NCR3/NKP30 or inflammatory factors such as CCL3/MIP1-α, LT (lymphotoxin) A/TNF-β, LTB/TNF-C. Conversely, these analyses did not evidence a strong up-regulation of well-expressed genes following TGF-β exposure. Most of the differentially expressed genes were related to either effector function, transcription, activation or metabolic functions (***Figure 3B***) (up-regulation, *red;* down-regulation, *blue*) upon TGF-β exposure. This cytokine diminished the expression levels of important genes related to effector functions such as CD40LG or CXCR6 described to regulate tissue trafficking of T lymphocytes (***Figure 3C***).

**Figure 3:**
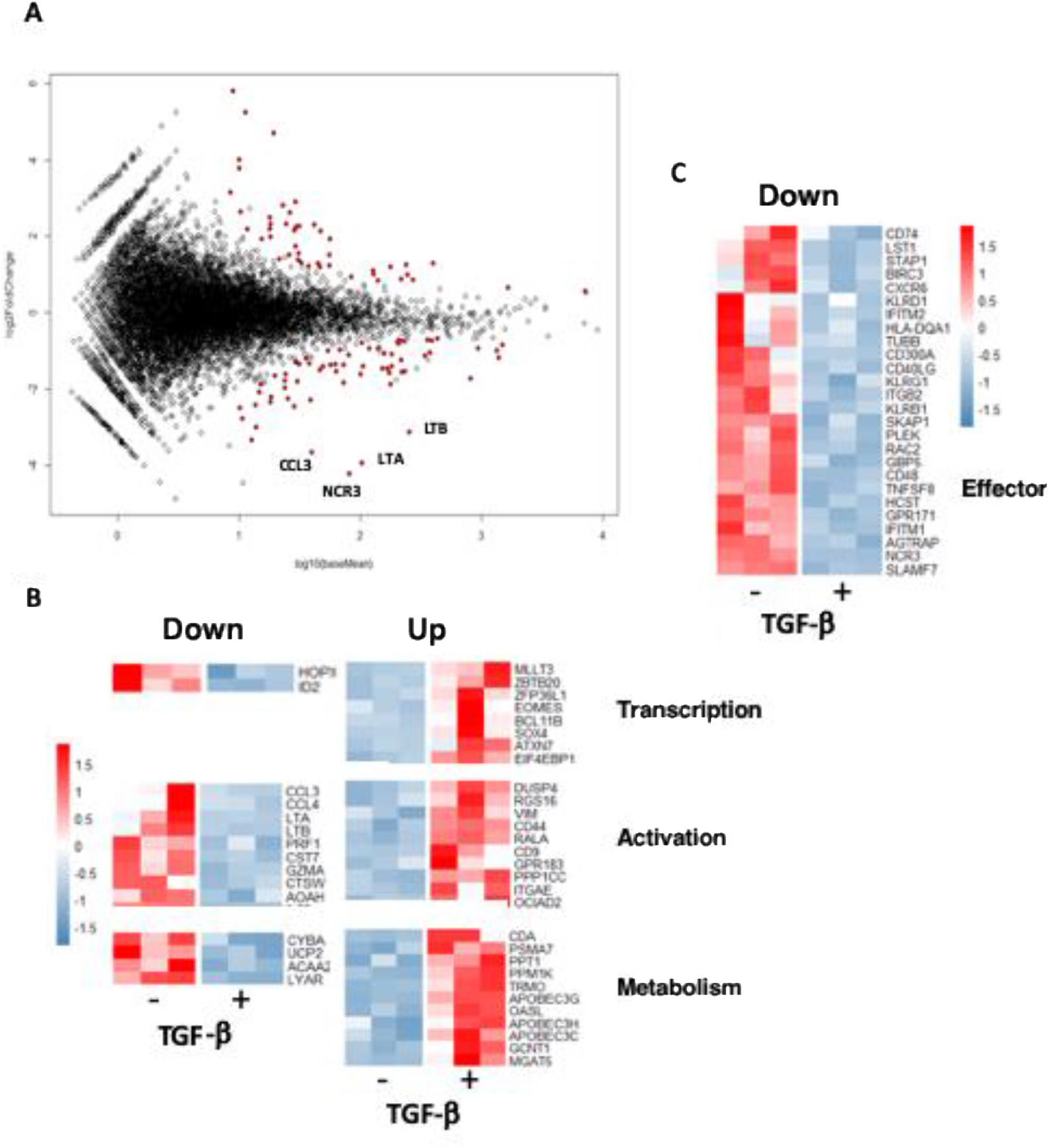
TGF-β exposure impacts the transcriptome of human Vγ9Vδ2 T cells. Human PBL-expanded Vγ9Vδ2 T cells were pretreated, or not, with TGF-β (10 ng/mL) for 3 days prior to RNA extraction. Comparative DGE-seq analysis was performed on bulk RNA-seq samples and data were analyzed using R software. MA plot **(A)** and heat maps **(B)** and **(C)** representing genes (indicated functions) that have been retrieved as differentially expressed following TGF-β exposure (+) or not (-). Colored scale, fold change expression. *Down*, down-regulated genes (*blue*); *Up*, upregulated genes (*red*).

These analyses show that the TGF-β induces the establishment of a particular transcriptional program in exposed Vγ9Vδ2 T cells which could be distinguished from regulatory or exhausted T cell signatures^17^.

### TGF-β alters the metabolic functions of Vγ9Vδ2 T cells

A growing number of studies shows that TME can alter the functions of infiltrating T cell effectors by affecting their metabolic activity^18,19^. To further investigate the existence of similar effects induced upon TGF-β exposure of human Vγ9Vδ2 T cells which was suggested by our transcriptome analyses, the energy metabolism was measured and compared in different conditions through real-time live cell analysis using Seahorse Extracellular Flux analyzer that enables direct quantification of mitochondrial respiration and glycolysis. γδ T cells were monitored at a resting state and after specific antigenic (BrHPP) or non-specific (PMA+ionomycin) activations (***Figure 4A***). In resting conditions, Vγ9Vδ2 T cells exposed to TGF-β (*closed symbols*) showed a weaker oxygen consumption (OCR, *oxygen consumption rate*) and media acidification rate (ECAR, *extracellular acidification rate*), as compared to their untreated counterparts. As expected, both specific and non-specific activations increased ECAR and OCR levels of control Vγ9Vδ2 T cells (*untreated*). However, a strong specific antigenic activation (BrHPP, 3 μM) of TGF-β-exposed Vγ9Vδ2 T cells induced barely detectable, if any, increase of ECAR or OCR metabolic activities. Moreover, upon exposure to TGF-β, OCR and ECAR levels were significantly lower in Vγ9Vδ2 T cells, as compared to untreated γδ T lymphocytes. Non-specific activation increased these metabolic levels without reducing these differences between these exposure conditions (***Figures 4B-D***). Moreover, TGF-β exposure reduces the viability of Vγ9Vδ2 T cells, as compared to untreated conditions (***Figures 4E-F)***. Taken together, the results show that TGF-β, which is produced in various TME of solid tumors, strongly impairs the metabolism of human Vγ9Vδ2 T cells by both reducing their mitochondrial respiration, their glycolytic activity and their viability which are key parameters for antitumor functions^20^.

**Figure 4.**
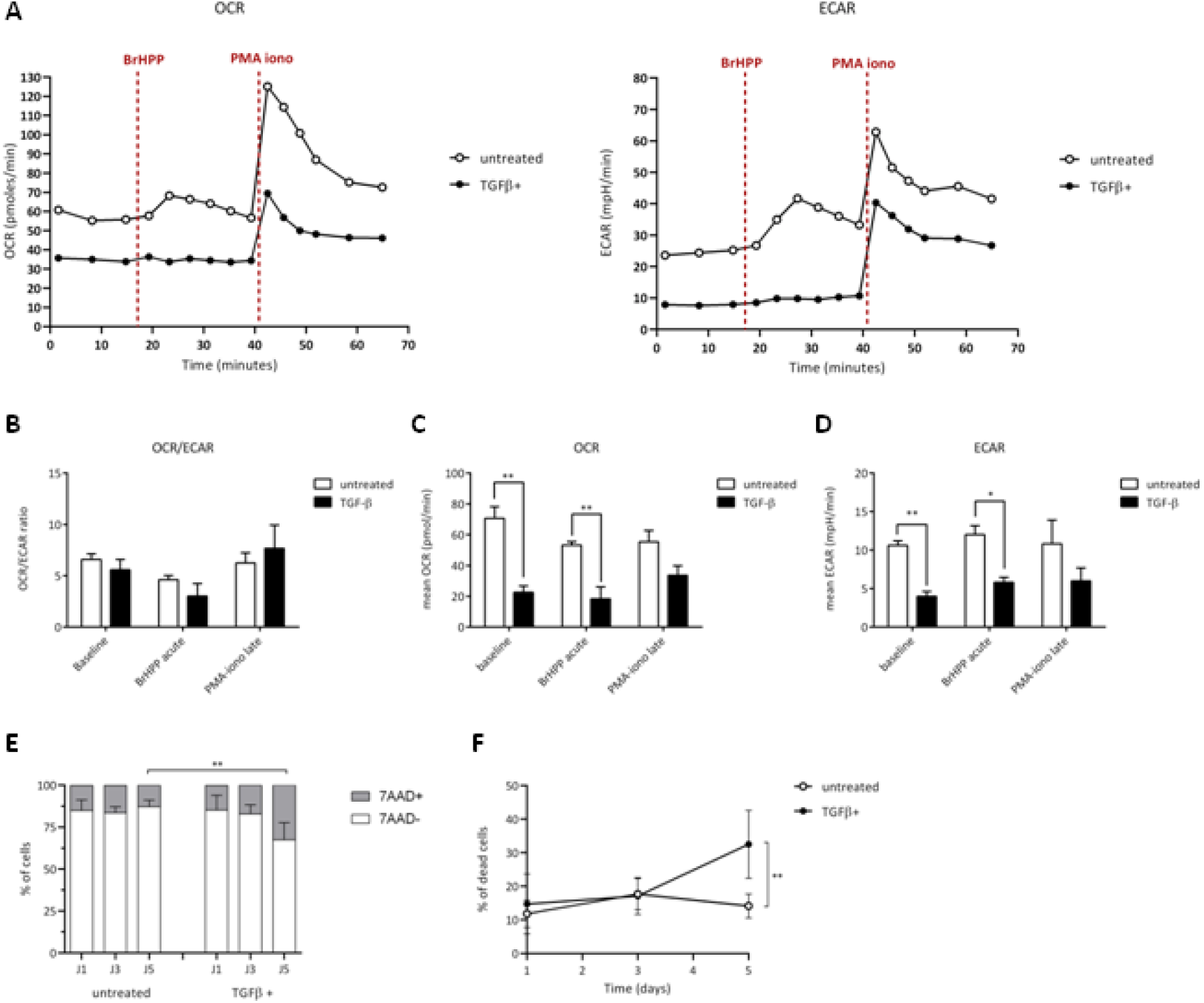
TGF-β alters the metabolism of Vγ9Vδ2 T cells. Vγ9Vδ2 T cells were pretreated (*closed symbols*) or not (*open symbols*) with TGF-β (10 ng/mL) for 3 days. Metabolic activities of T lymphocytes were measured using Seahorse XF8. **(A-D)** Monitoring of mitochondrial oxygen consumption rate (OCR) and extracellular acidification rate (ECAR) were performed at resting state, after BrHPP stimulation or PMA-ionomycin injection as described in the *Materials and Methods* section. **(A)** Represensative experiments. Rates were calculated as follows: difference between the mean of the 3 values in the absence of substrate and the mean of the 4 values after injection of the substrate, as well as a ratio of the 2 for each condition **(B-D). (E)** and **(F)** Cell death induced by TGF-β added within the media of Vγ9Vδ2 T cells (10 ng/mL for 3 days) was measured by measuring fluorescent non-viable cells following 7-AAD staining after the indicated culture timepoints days of culture. n=6, mean ± SD. Mann-Whitney ^*^p<0,05; ^**^p<0,01. PMA-iono: PMA + inomycin (positive control).

## Discussion

Our results indicate that TGF-β, which transcription is detected at high levels in brain tumors with a poor prognosis, interferes with the antigenic activation of human Vγ9Vδ2 T cells *in vitro*. Importantly, they show that this tumor-produced regulatory cytokine strongly impairs their antitumor cytolytic reactivity, through multiple phenotypic, transcriptomic changes as well as metabolic alterations.

Regarding tumor surveillance, the immune system orchestrates with broad molecular and cellular networks for sensing and eliminating tumor cells. Numerous studies focused on innate-*like* T cell subsets including γδ T cells, and particularly on non-alloreactive peripheral human Vγ9Vδ2 T cells which represent promising effector candidates for designing novel immunotherapies. The implication of Vγ9Vδ2 T cells in antitumor immune response is supported by the identification of tumor-infiltrated γδ T cells, which might be considered as a a prognosis tool for some oncological indications^21,22^. However, the clinical efficacy of Vγ9Vδ2 T cell-based therapies is still disappointing. Although their conserved and MHC-independent specificity is expected to overcome issues related to antigen presentation and recognition, their efficacy against solid tumors remains limited and hampered by various immunological, biochemical and tumor-linked constraints. Once T cells have successfully reached and infiltrated the tumors, they need to evolve within a strong immunoregulatory environment that aims at both taming their antitumor functions and promoting tumor development.

TME is a complex network constituted by cancer-associated fibroblasts, myofibroblasts, extracellular matrix, immune/inflammatory cells, blood, and vascular networks as well as signaling components including cytokines, chemokines and growth factors. Previous studies have shown the sensitivity of Vγ9Vδ2 T cells to two TME-produced immunosuppressive factors: PGE2 and TGF-β^23,24^. PGE2 potently suppresses the proliferation, cytokine production, and cytotoxic capacity of γδ T cells. TGF-β, a secreted dimeric multifunctional cytokine that signals via specific receptors on plasma membranes, contributes to cancer progression by its ability to promote epithelial-to-mesenchymal transition of cancer cells, angiogenesis, and immune evasion^25^. We showed that TGF-β exposure severely impacts the cytolytic activity of human Vγ9Vδ2 T cells. Interestingly, this inhibition was durable *in vitro* (up to 3 days) and reversible in the absence of the cytokine. TGF-β exposure reduces the expression of lytic molecules (e.g. perforin, granzyme B), as well as surface molecules mediating this antitumor reactivity such as NKG2D, as described for NK and CD8^+^ T cells^12^. NKG2D was extensively described as a key receptor for sustaining the reactivity of various lymphoid subsets, including human Vγ9Vδ2 T cells, against several tumor cells. Supporting this observation, we previously evidenced the major role played by NKG2D for the elimination of human mesenchymal GBM tumor cells by allogeneic human Vγ9Vδ2 T cells^16^ which can infiltrate GBM^26^. Therefore, in such pathological contexts, one could expect that NKG2D-controled antitumor Vγ9Vδ2 T T cell reactivities would be altered by TGF-β whose transcription is elevated in brain tumors^27,28^ as previously described for NK and CD8^+^ T cells^29^.

Studies reported opposite effects of this cytokine in which the antitumor activity of TGF-β-conditioned Vγ9Vδ2 T cells is potentiated and could be used for improving their efficacy in adoptive transfer therapies^30,31^. This highlights the complex pleiotropic effects of some regulatory cytokines on effector T cells, according to the context of exposure (e.g. sensitization of resting T cells *vs* conditioning for amplification) which might introduce several different parameters such as the cellular environment, the kinetics as well as the differentiation status/plasticity of T cells. As shown here, TGF-β does not only directly inhibit the antitumor reactivity of infiltrated Vγ9Vδ2 T cells but also induces specific modifications of their transcriptional program and phenotype. Comparative RNA-seq analysis performed on resting Vγ9Vδ2 T cells exposed to TGF-β, did not show any polarization towards Th17 (e.g. STAT3, IL-22), Th9 (e.g. IRF4, IL-9) or regulatory (e.g. FoxP3) functional profiles. The lack of TGF-β-induced regulatory functions was also confirmed in T cell proliferation assays *in vitro* (*data not shown*). TGF-β exposure affects the expression of genes either implicated in lymphocyte activation, transcription effector processes (downregulation). Interestingly, as already observed for NK cells^32^, the expression of NKG2D as well as NCR3/NKp30 is strongly impacted by TGF-β exposure. We showed that this cytokine impacts particular metabolic pathways in human Vγ9Vδ2 T cells. Both glycolysis and mitochondrial respiration, which are key metabolic anti-tumor effector pathways^33^ are strongly abated upon cytokine exposure. This plunges TGF-β-exposed Vγ9Vδ2 T cells in a hypoxia*-like*-induced state through raising their activation threshold, thus leading to a dramatic reduced reactivity against tumor cells.

Altogether, the results indicate that TGF-β, a cytokine that is massively produced in the TME of various solid tumors, acts as a potent taming cytokine on resting PBL-expanded human Vγ9Vδ2 T cells, at different levels. It does not only directly target the key NKG2D axis, to inhibit the antitumor cytolytic activity but also affects their transcriptional and metabolic program to reduce their viability and functional activity. This should be taken in account for designing improved Vγ9Vδ2-based cancer immunotherapies, by proposing genome editing technologies (e.g. CRISPR-Cas9) for preparing engineered Vγ9Vδ2 T cells that can effectively counter tumor escape promoted by TME-derived factors^34^.

## Acknowledgments

We thank the Flow Cytometry and FACS core facility “Cytocell” (BioCore, SFR Bonamy, Biogenouest, LabExIGO, Nantes, France) for their expert technical assistance, B. Navet and the BIRD facility for expert bioinformatic assistance.

## Disclosure of interest statement

The authors report there are no competing interests to declare.

## Data availability statement

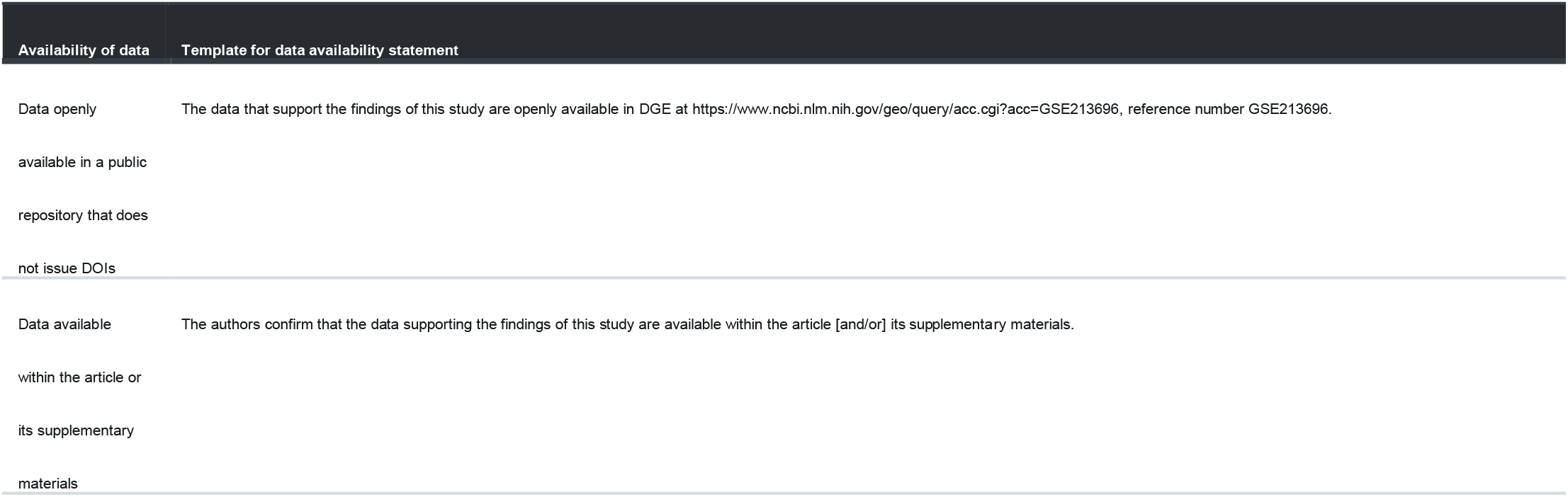

## Notes

### Competing Interest Statement

The authors have declared no competing interest.

