## Supplemental Figure 1 for "A dramatic impairment of the antitumor activity of human Vγ9Vδ2 T cells is induced by TGF-β through significant phenotype, transcriptomic and metabolic changes"

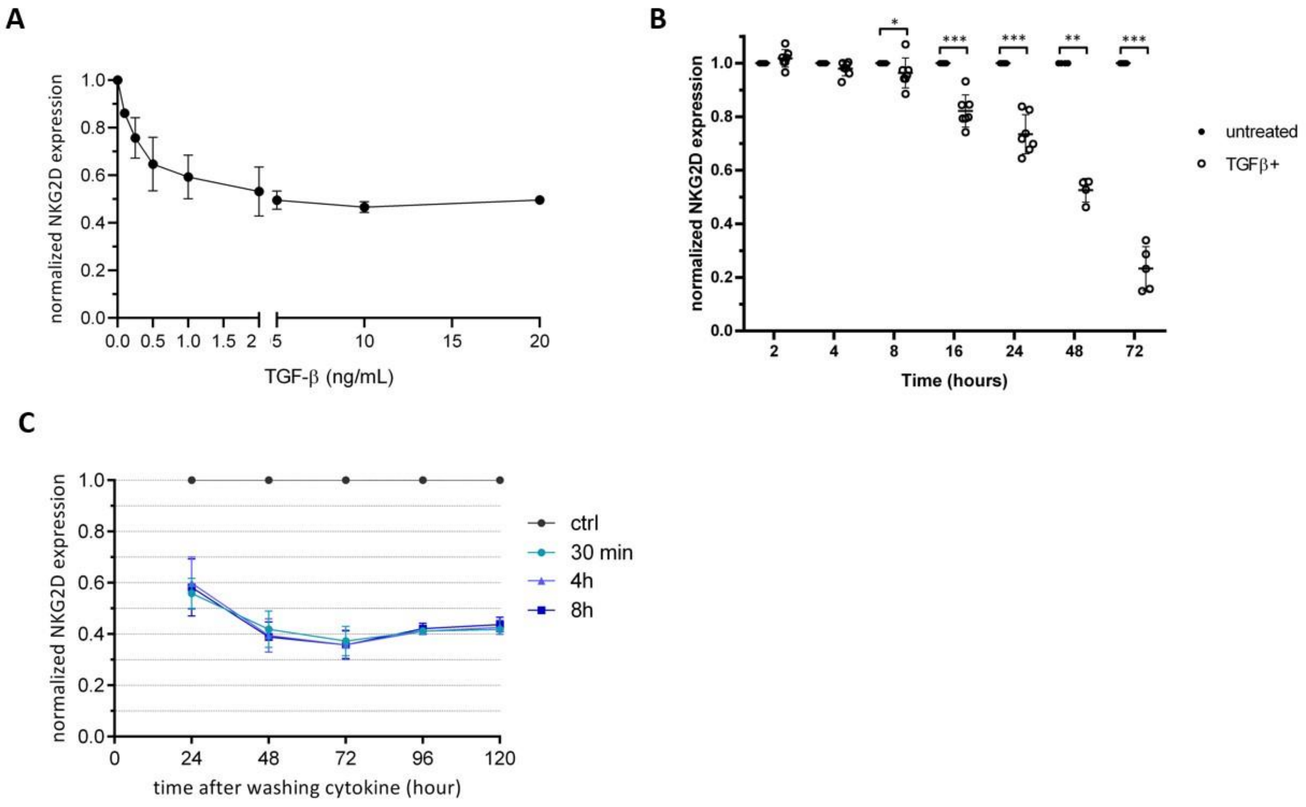


**Supplemental Figure S1.** **(A)** Normalized expression of NKG2D measured by flow cytometry at the surface of human Vγ9Vδ2 T cells incubated for 3 days with indicated increasing doses of recombinant human TGF-β. **(B)** Kinetics of expression of NKG2D expressed at the surface of human Vγ9Vδ2 T cells at different time-points, following addition of TGF-β (10 ng/mL). Cell surface stainings were measured by flow cytometry. **(C)** NKG2D expression at the surface of human Vγ9Vδ2 T cells incubated with recombinant human TGF-β (10 ng/mL) at the indicated time-points (ctrl, negative control). NKG2D expression levels were next measured and normalized, at the indicated time-points after washing the cells. n>5 Mann-Whitney tests *p<0.05; **p<0.01.
